# Using Experimental Evolution to Correct Mother-Daughter Separation Defects in Brewing Yeast

**DOI:** 10.1101/2025.11.25.687580

**Authors:** Lauren M Ackermann, Amanda Ro, Barbara Dunn, Joseph O Armstrong, Ryan Moore, Greg Doss, Maitreya J Dunham

**Author notes:** Contributed equally to this work (Lauren M Ackermann, Amanda Ro).

## Abstract

The budding yeast, *Saccharomyces cerevisiae*, is the workhorse of the brewing industry. Brewers have domesticated a vast array of different strains with traits that complement the beers they wish to brew. Yet, some domesticated strains also harbor traits that are undesirable. One example of an undesirable trait in brewing strains is the mother-daughter separation defect (MDSD). MDSDs are a reproductive flaw present in a widely used brewing strain, London Ale III. MDSDs cause cells to form large clusters, possibly leading to the known requirement for more headspace during London Ale III fermentations that result in a lower fermentative yield. Because MDSDs can be caused by mutations to a number of genes, targeted genetic approaches to reduce MDSDs are experimentally challenging, especially for a tetraploid strain like London Ale III. To improve MDSDs in this strain, we employed experimental evolution and passaged populations from three biological replicates for over 200 generations to generate three independent evolved strains that form fewer clusters than the ancestral strain, as seen by microscopy. To confirm these results, we used flow cytometry to measure the average size of the clusters in clones of our evolved replicates and found them to be smaller on average than the ancestor. We also qualitatively assessed the aggregation phenotype using a settling assay and found that our evolved replicates settle slower than the ancestor. Finally, we characterized the mutations in our evolved replicates using whole genome sequencing and identified increased copy numbers of chromosome 1 and chromosome 14 in all three evolved clones. The best-performing strain generated by this project is now available commercially. This project demonstrates how experimental evolution can be used to select against less desirable traits in industrial yeast strains when targeted genetic approaches present considerable challenges. Future research could implement a similar approach to improve other traits in widely used brewing and baking strains.

## Description

*Saccharomyces cerevisiae* replicate through a process known as budding, where a mother cell undergoes cytokinesis to form a daughter cell, which emerges as a bud. During this process, the cytoplasm is divided by the contraction of the actomyosin ring and by the formation of a specialized wall between the mother and bud called the septum. Cell division is successful when the septum is completely broken down, allowing the mother and daughter cells to fully separate (Weiss 2012). When the septum is unable to fully break down, mother and daughter cells remain connected leading to a mother daughter separation defect (MDSD). MDSDs cause cells to form large chains or clusters that are joined together at bud necks. These clustered cells are sometimes referred to as “snowflake yeast” due to their snowflake-like appearance (Ratcliff et al. 2012). Brewing strains with MDSDs accumulate large amounts of floating foam during active fermentation, requiring specialized release valves and/or more starting headspace (less wort) in the fermenter, resulting in lower fermentative yields. It is possible that the “snowflake” cell clusters could be responsible for this behavior. Despite this disadvantage, many brewers still choose to use strains with MDSDs due to other desirable traits these strains hold. The goal of this project was to correct MDSDs in a widely used and commercially available brewing strain, London Ale III, the most common strain used to make New England style (Hazy) IPAs. Correcting MDSDs in a targeted way is challenging because MDSDs can be caused by mutations to multiple genes, for example *AMN1* (Fang et al. 2018) and *ACE2* (Voth et al. 2005, Ratcliff et al. 2015). Moreover, London Ale III is tetraploid which further complicates a targeted genetic approach. To overcome this challenge, we sought to ameliorate MDSD in the London Ale III strain by using experimental evolution to select against aggregating cells, as previously described (Amorosi 2020, Ratcliff et al. 2012).

In laboratory culture, large, clustered aggregates are less buoyant and settle faster than single cells. In our experimental evolution protocol, we took advantage of this property by using gentle centrifugation (100 x g) for a short duration (20 seconds) to enrich smaller clusters and single cells in the supernatant. We then transferred the supernatant into fresh media to complete the cycle (Fig. 1A). We repeated this cycle 37 times (approximately 200 generations) for three biological replicates, then isolated clones from each of the three evolved populations to generate novel strains (YMD5227, YMD5228, and YMD5229). We observed by microscopy that large cell clusters decreased in frequency in all three of our evolved clones, while the frequency of single cells and smaller clusters increased compared to the ancestor (representative image in Fig. 1B). We confirmed these observations both qualitatively and quantitatively with three orthogonal measurements.

**Figure 1.**
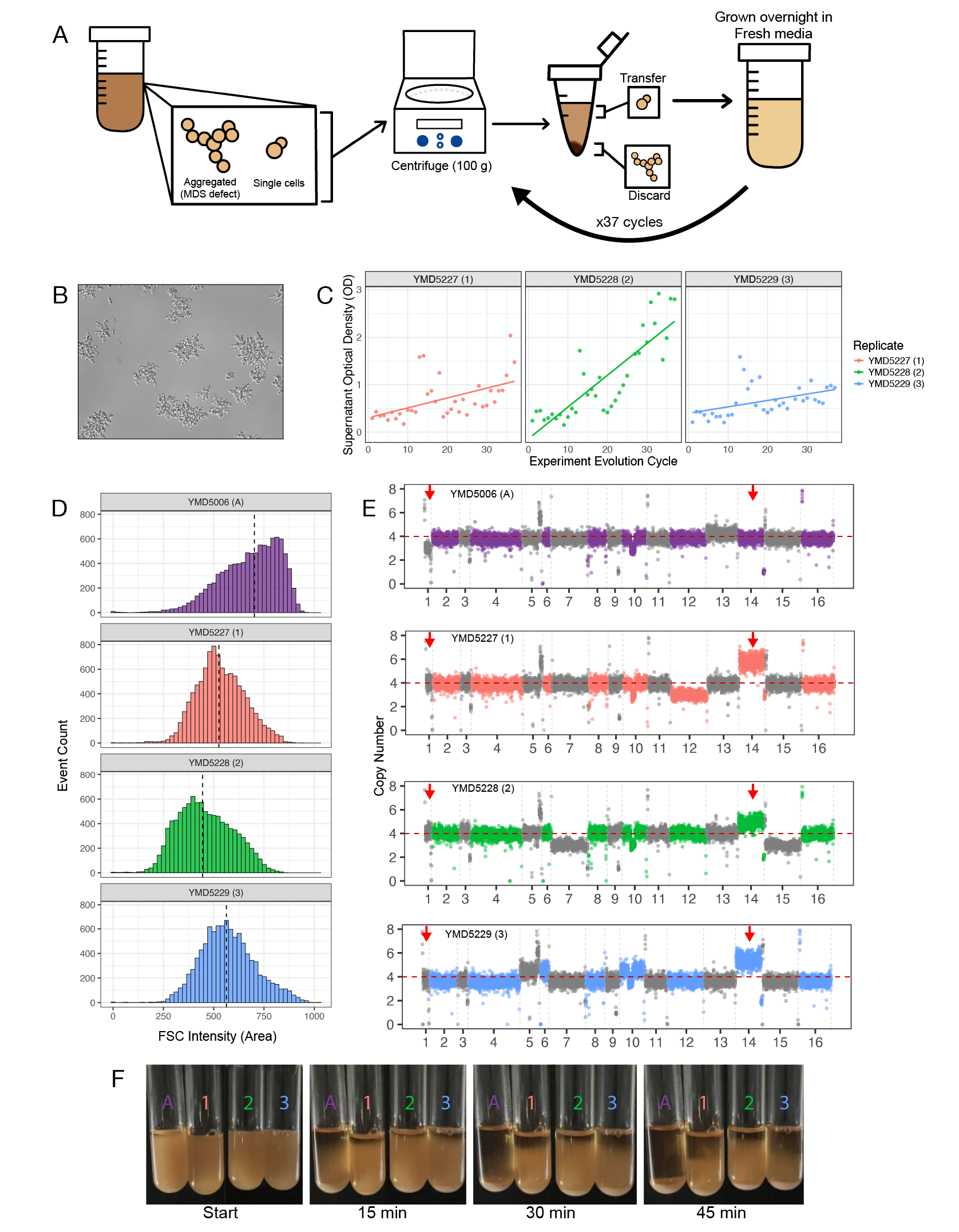
Improving MDSDs with experimental evolution. **A**. A schematic of our experimental evolution cycle, performed daily with three replicate populations. **B**. Microscope image at 20x magnification of an overnight culture from the ancestral strain. **C**. Plot showing supernatant optical density (OD600) over time, measured by experimental evolution cycles. Each replicate is indicated by a different color: red, green, and blue representing the 1st, 2nd, and 3rd replicate, respectively. A line of best fit in the same color is plotted to demonstrate the general trend in optical density. **D**. Histograms demonstrating forward scatter (FSC, a proxy for cell and cluster size) of an individual clone isolated from the ancestor and each replicate, measured with flow cytometry. Event count is shown on the y-axis, while FSC-A is shown on the x-axis. The median cell/cluster size is approximated and indicated by the dashed black line on each histogram. Purple, red, green, and blue correspond to the ancestor, replicate 1, replicate 2, and replicate 3, respectively. **E**. Copy number plots of the ancestor clone, clone from 1st replicate (YMD5227), clone from 2nd replicate (YMD5228), and clone from 3rd replicate (YMD5229) generated with Illumina sequencing using the read depth across 1000 base pair rolling windows. **F**. Qualitative settling assay demonstrating settling rates over time of three replicate clones. Culture tubes are shown in 15-minute intervals and different colors represent the ancestor clone and three replicate clones.

First, we measured the optical density of the supernatant (Fig. 1C) during each experimental cycle and found that it increased over time in all three of our replicates, suggesting a higher proportion of single and small clusters. After 23 evolution cycles, we began taking optical density measurements of the overnight culture before centrifugation. We found that over time, the ratio of supernatant OD to overnight culture OD increased, validating that single and small clusters increased in our replicates. We also conducted a qualitative settling assay, where tubes containing overnight cultures of our three evolved clones remained turbid for longer than an overnight culture of the ancestral strain. After 45 minutes, we found the ancestral strain completely settled, while the evolved clones remained cloudy (Fig. 1F). We performed this assay with our evolved replicate populations at three other timepoints during the evolution. All settling assays confirmed that our evolved populations settled notably slower than the ancestor. Finally, we quantitatively assessed the size distribution of the evolved clones (YMD5227, YMD5228, and YMD5229) using flow cytometry to measure the intensity of forward light scattering, which correlates with cell or cluster size. We generated histograms of forward scatter intensity for the three evolved strains and compared them with the ancestor strain. All three evolved strains had a lower median forward scatter than the ancestor (Fig. 1D), indicating that the evolved strains form smaller clusters and contain more single cells than the ancestor. These results indicate that we successfully selected against cells with MDSDs and generated strains that feature more single cells and fewer large clusters. We identified from our settling assay and cytometric analysis that the evolved strain YMD5228 settled the slowest and had the smallest average cluster size.

To uncover the genetic basis of the improvement in MDSD in our evolved clones, we performed whole genome sequencing on the evolved strains and the ancestral strain. We analyzed the sequences of the evolved strains and searched for single nucleotide polymorphisms (SNPs) or copy number variation that may explain the improved MDSD in our evolved strains. We first looked specifically for mutations in *ACE2, AMN1, CTS1, DSE4, DSE2, SUN4, DSE1, IZH4*, and *SCW11*, genes known to be associated with the snowflake phenotype (Ratcliff et al. 2015). Across all evolved strains, we did not find new SNPs in these genes, only changes to existing allele frequencies (see below). We then looked at genes involved in *AMN1* negative selection, *STE11, RIM8, RIM13, RIM20*, and *RIM101* (Sniegowski 2025). Again, we did not find meaningful and novel SNPs in our evolved clones, only to preexisting allele frequencies. We also did not find evidence for loss of heterozygosity, a hallmark of mitotic recombination (Zhu et al. 2025).

To explore whether changes to whole chromosome copy numbers could account for the improvement in MDSD, we generated copy number plots of the ancestor and evolved strains using the read depth across 1000 base pair rolling windows (Fig. 1E). We found that each evolved strain experienced a copy number increase of chromosome 14, as well as of chromosome 1. Chromosome 1 increases from three copies in the ancestral strain to four copies in all of the evolved strains. Chromosome 14 increases from four copies in the ancestral strain to five copies in YMD5228 and six copies in YMD5227 and YMD5229. Allele frequencies of SNPs in chromosome 1 were consistent with three copies for the ancestor, but four copies for the evolved clones. We saw similar patterns in allele frequencies of SNPs located on chromosome 14, consistent with six copies in YMD5227 and YMD5229, and five copies in YMD5228. Different copies of each chromosome were amplified in each clone. We observed other abnormal chromosome copy numbers–or aneuploidies–in the evolved strains; however, only the increases in copy number of chromosome 1 and 14 are shared between all three evolved strains. Two of our candidate genes, *DSE4* and *SUN4*, are located on chromosome 14. Thus, the increase in copy number to either (or both) chromosome 1 and 14 may play a role in improving MDSD in this strain.

Using experimental evolution, we successfully corrected the MDSD in London Ale III. The strain YMD5228, which showed the most improvement to MDSD, is now commercially available to brewers. In the future, we want to identify specific causal genes that were involved in driving a non-aggregation phenotype in these evolved strains. We hope to identify which genes on chromosomes 1 and 14 might affect the phenotype, enabling a more targeted approach to engineering other strains. This could provide greater insight into abnormal yeast cell separation and the genotypic changes driving this phenotype in a tetraploid brewing strain.

## Methods

### Strains/Media

10% malt extract media was made by adding 50g of dry malt extract (Briess CBW Pilsen Light Dry Malt Extract #5760) to a sterile jug, then topping off with distilled water to a final volume of 500mL. This mixture was autoclaved then decanted to remove large particulates. Finally, media was vacuum filtered to remove any remaining particles and ensure sterility.

### Experimental Evolution Cycle

We performed a modified method to one previously described (Amorosi 2020, Ratcliff et al. 2012). Overnight cultures were vortexed and 1.5mL was transferred to a sterile Eppendorf tube. Tubes were briefly vortexed, then centrifuged at 100 x g for twenty seconds. 1mL of supernatant was immediately pipetted into culture tubes containing 4mL of 10% malt extract (see Fig. 1A). Pipetting 1mL of supernatant was done carefully by moving down from the top of the tube in order to capture the smallest clusters and avoid the pellet. Culture tubes were then vortexed and placed on a roller drum at 30ºC to grow overnight. Cells were propagated daily in this manner for 37 evolution cycles (approximately 200 generations).

### Supernatant Optical Density Measurements

Optical density (OD600) measurements were taken using BioRad SmartSpec™ 3000. Overnight cultures of each replicate were vortexed, then were transferred to a 1.5mL eppendorf tube. Tubes were vortexed, then centrifuged at 100 x g for twenty seconds. 1mL of the supernatant was pipetted by carefully moving down from the top of the tube then transferred into a clean cuvette. The spectrophotometer was blanked using 1mL of 10% malt extract. After 23 evolution cycles, optical density measurements were also taken of the overnight culture in a similar manner. Mixtures were diluted as needed to remain in the linear range of the spectrophotometer (<1.0 OD).

### Generation Number Estimation

Generations were defined as a doubling of the estimated cell mass, as measured by optical density. The generations between each cycle were estimated using OD600 of the transferred supernatant and the overnight culture using the following equation:

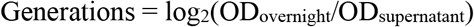

### Settling Assay

Culture tubes containing the evolved replicates and the ancestor were thoroughly vortexed, clamped onto a lab stand, and imaged every 15 minutes for a total of 90 minutes. Images were taken with an iPhone 13 clamped at the same position over the course of the assay.

### Flow Cytometry

Overnight cultures of each strain were diluted 1:100 and forward scatter intensity was measured on a Sony SA3800 flow cytometer. At least 10,000 events were recorded per sample.

### Illumina sequencing

Genomic DNA was isolated from the ancestor and a single clone from each of the three evolved populations using Hoffman/Winston phenol chloroform extraction (Hoffman et al. 1987). Genomic DNA was purified using the Zymo Research DNA Clean & Concentrator-5. The purified DNA was then prepared for Illumina short read sequencing using the Nextera XT DNA Library Preparation Kit. Each sample received >10 million paired end reads. Determination of indels, copy number variation, and SNPs followed a modified protocol as previously described (Taylor et al. 2022). Short read sequences were aligned to the *Saccharomyces cerevisiae* reference genome (SacCer3) then copy number plots were generated by analyzing read depth across 1,000 base pair sliding windows, normalizing to read depth across all chromosomes, and normalizing to the ploidy of the strain (Large et al. 2020). Sequencing data is available in the NCBI Sequence Read Archive (SRA) under BioProject PRJNA1358295.

## Reagents

| Strain | Species | Source/Reference |
| --- | --- | --- |
| Juice (London Ale III) | *Saccharomyces cerevisiae* | Imperial Yeast |

## Acknowledgments

We thank Owen Lingley for advice and support. We thank the rest of the Dunham lab for advice on this project and related troubleshooting.

## Author Contributions

Lauren M Ackermann: Data curation, Investigation, Formal analysis, Visualization, Writing - original draft, Writing - review & editing

Amanda Ro: Data curation, Investigation, Formal analysis, Visualization, Validation, Writing - original draft, Writing - review & editing

Barbara Dunn: Conceptualization, Project administration, Writing - review & editing

Joseph O Armstrong: Investigation, Formal analysis, Visualization, Writing - review & editing Ryan Moore: Conceptualization, Investigation, Writing - review & editing

Greg Doss: Conceptualization, Investigation

Maitreya J Dunham: Conceptualization, Funding acquisition, Project administration, Resources, Supervision, Validation, Writing - review & editing

